# Target-dependent modulation of finely tuned gamma by transcranial ultrasound in Parkinson’s disease

**DOI:** 10.64898/2026.11.26.769557

**Authors:** A. Asadian, C. Sarica, H. Ramezanpour, D. Sastin, R. Annirood, A. Bhattacharya, J. Yuen, A. Loh, F. Villamil, J. Cabrera-Montes, C. Chow, F. A. Schmidt, J. F. Nankoo, R. P. Munhoz, T. A. Valiante, A. Fasano, M. Hodaie, S. K. Kalia, R. Chen, A. M. Lozano, G. Darmani

## Abstract

**Background:** Finely tuned gamma (FTG) is a narrowband oscillation associated with dopaminergic ON states and dyskinesia in Parkinson’s disease (PD), although its physiological significance remains debated. While deep brain stimulation (DBS) can entrain gamma oscillations, whether noninvasive ultrasound can modulate endogenous FTG across motor circuits remains unclear.

**Methods:** We analyzed local field potentials recorded using sensing-enabled DBS systems in two cohorts of patients: 17 with subthalamic nucleus (STN) and 10 with globus pallidus internus (GPi) electrodes. Theta-burst transcranial ultrasound stimulation (tbTUS) targeted the primary motor cortex (M1) or GPi. FTG was assessed before, during, and up to 45 minutes after sonication. Associations with beta activity, levodopa-equivalent daily dose (LEDD), and dyskinesia were also examined.

**Results:** FTG was detected in eight STN patients and one GPi patient, with five exhibiting reproducible peaks suitable for longitudinal analysis, broadly consistent with previous reports of intermittent FTG expression. M1 sonication reduced STN FTG in three patients, whereas GPi sonication increased STN FTG in one but did not significantly alter locally recorded GPi FTG in another. Finger-tapping-related FTG responses were heterogeneous. Resting FTG was not significantly correlated with beta activity or LEDD. However, dyskinesia was more frequent in sessions with detectable FTG than in those without (47% vs. 20%), although this association was not statistically significant (p = 0.067).

**Conclusions:** These findings suggest that noninvasive ultrasound can modulate endogenous FTG in a circuit-dependent manner, distinct from DBS-induced gamma entrainment. FTG may provide a complementary electrophysiological biomarker alongside beta oscillations, potentially capturing additional aspects of motor-network activity and dyskinesia. Combining noninvasive TUS with intracranial DBS sensing may offer a useful approach for investigating FTG and its potential application in future adaptive DBS and individualized neuromodulation strategies.

## 1. Introduction

Finely tuned gamma (FTG) is a narrowband oscillation, typically centered around 65–90 Hz [1, 2], in which the physiological significance in Parkinson’s disease (PD) remains incompletely understood. Initially associated with levodopa-induced dyskinesia, FTG was identified in the motor cortex and subthalamic nucleus (STN), with increased cortico-subthalamic coherence during dyskinetic states [2]. Subsequent chronic recordings further supported an association between narrowband gamma activity and dyskinesia [1]. However, more recent evidence has challenged the specificity of FTG as a dyskinesia biomarker [3–5]. Ambulatory recordings demonstrated that FTG tracks dopaminergic medication cycles independent of dyskinesia [4], while a larger electrophysiological study showed that spontaneous and stimulation-entrained FTG can occur during therapeutic ON states without accompanying dyskinesia [5]. These findings suggest that FTG may reflect a broader physiological state associated with dopaminergic modulation and motor network activity, rather than exclusively representing pathological hyperkinesia [4–6].

Beyond its clinical associations, FTG has attracted increasing interest as an electrophysiological marker of basal ganglia network dynamics. Unlike the broader gamma activity associated with movement and sensorimotor processing [7], FTG appears as a relatively narrow spectral peak, suggesting the involvement of synchronized neuronal populations or resonant network activity [6, 8]. Its expression is heterogeneous across patients and recording sites, and can vary with medication, movement, and stimulation state [3–5, 9, 10]. Levodopa-induced STN FTG has been observed during voluntary movement, with transient increases in amplitude around movement onset [10]. Repetitive finger tapping has also been shown to temporarily enhance DBS-entrained gamma activity, with entrainment associated with improved motor performance [11]. However, whether endogenous FTG is consistently modulated during motor tasks, and how these responses are affected by neuromodulation, remains unclear. Understanding how FTG is generated and modulated across different basal ganglia circuits and behavioural states is therefore important for interpreting its potential role as an electrophysiological biomarker.

Deep brain stimulation (DBS) provides further evidence that FTG is sensitive to external modulation of motor circuits. High-frequency stimulation of STN or GPi can entrain gamma oscillations at approximately half the stimulation frequency, an effect associated with improved motor performance and altered relationships between spontaneous gamma activity and dyskinesia [2, 11, 12]. More recently, subharmonic gamma entrainment during thalamic DBS for multiple sclerosis tremor has suggested that the capacity to generate and entrain narrowband gamma extends beyond the parkinsonian basal ganglia [13, 14]. Nevertheless, stimulation-induced entrainment and modulation of spontaneous FTG may reflect distinct physiological mechanisms [3, 12]. Whether changes in FTG represent local responses to stimulation or broader alterations in network interactions remains unresolved, particularly across different cortical and subcortical targets.

Transcranial ultrasound stimulation (TUS) is an emerging noninvasive neuromodulation technique capable of targeting both cortical and deep brain regions with relatively high spatial precision [15]. Recent studies combining TUS with intracranial local field potential (LFP) recordings have demonstrated target- and state-dependent modulation of basal ganglia oscillations in PD [16, 17]. These findings establish TUS as a promising approach for investigating how perturbations of distinct motor-network nodes influence activity in interconnected deep brain structures. This approach may be particularly informative for FTG, whose sensitivity to stimulation location and relationship to distributed motor circuits remain poorly characterized.

Here, we investigated FTG modulation during theta-burst TUS (tbTUS) of the primary motor cortex (M1) or globus pallidus internus (GPi), using LFPs recorded from implanted sensing-enabled DBS leads in two independent cohorts: 17 patients with STN leads and 10 patients with GPi leads. We examined whether endogenous FTG could be modulated by noninvasive ultrasound and whether its response differed according to the stimulation target, recording site, and behavioural states.

## 2. Materials and methods

### 2.1. Study cohorts and participant selection

This analysis used intracranial LFP recordings from two previously reported cohorts undergoing TUS with sensing-enabled DBS leads [16, 17]. The first cohort comprised 17 patients with STN leads, whereas the second included 10 patients with GPi leads. Both studies previously demonstrated electrophysiological modulation of basal ganglia activity following TUS [16, 17]. Both parent studies received approval from the University Health Network Research Ethics Board. All patients provided written informed consent before participation, and the studies were conducted in accordance with the Declaration of Helsinki [16, 17].

### 2.2. Electrode localization and anatomical visualization

For the STN cohort, contact coordinates in MNI space were taken from [16]. There, leads were localized with Lead-DBS v3.0 from pre- and postoperative T1-weighted MRI, normalized to MNI ICBM 2009b nonlinear asymmetric space using ANTs SyN [18] with subcortical refinement, and reconstructed semi-automatically with manual adjustment. For the GPi cohort, a similar process was followed with electrode segmentation performed in Brainlab (Brainlab SE, Munich, Germany). The BigBrain 300 μm volume [19] and the Pauli probabilistic subcortical nuclei atlas [20] were transformed into MNI template space. Three-dimensional rendering was performed using PyVista [21]. Localized DBS leads for both cohorts are shown in Fig. 1A.

**Figure 1.**
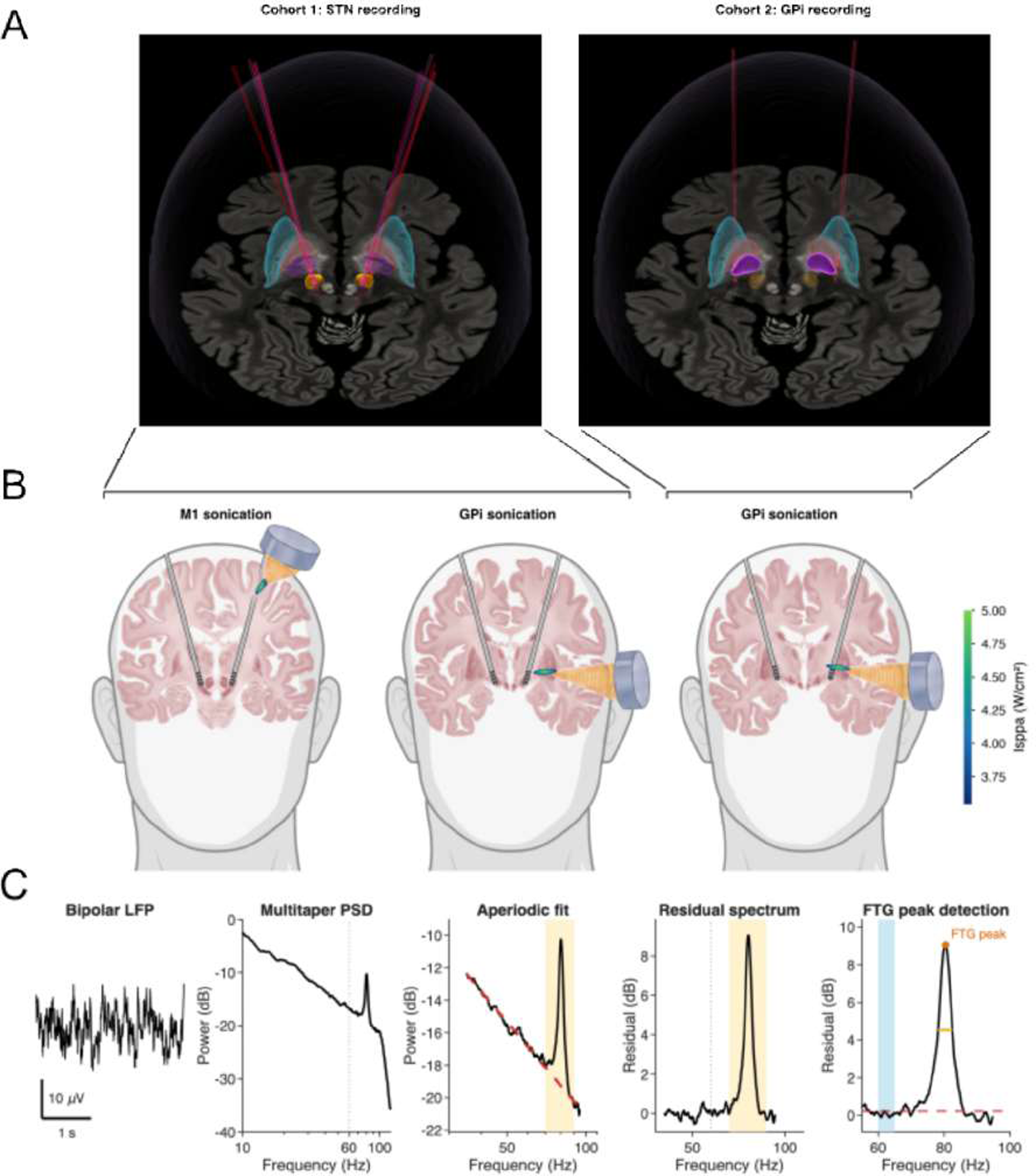
Recording sites, ultrasound targets, and FTG detection. (A) Reconstructed DBS electrode trajectories for Cohort 1 (STN recording, n = 17; left) and Cohort 2 (GPi recording, n = 10; right), shown on a coronal BigBrain template section [19] with the putamen (cyan), GPe (orange), GPi (magenta) and STN (yellow) from the Pauli atlas [20]. The recorded nucleus is highlighted in each panel. (B) Simulated 500-kHz acoustic fields on coronal BigBrain template [19] sections illustrating M1 sonication with STN recording (left, n = 3), GPi sonication with STN recording (middle, n = 1), and GPi sonication with GPi recording (right, n = 1). Colour scale indicates estimated in situ Iₛₚₚₐ. (C) Representative FTG detection, showing the bipolar LFP, multitaper spectrum, aperiodic fit, and residual spectrum with a detected peak. FTG was identified within 70–90 Hz using a threshold derived from the 60–65 Hz reference band.

### 2.3. Ultrasound delivery and recording epochs

TUS was delivered using a 500-kHz ultrasound system with individualized targeting, as described in the parent studies [16, 17]. Both cohorts used the NeuroFUS transducer power output system equipped with a four-element annular array transducer (CTX-500; Brainbox Ltd., Cardiff, UK). The present analysis included theta-burst TUS (tbTUS), consisting of 20-ms pulses repeated every 200 ms (5 Hz pulse repetition frequency, 10% duty cycle) for 120 s per hemisphere. In both cohorts, the spatial-peak pulse-average intensity (Iₛₚₚₐ) was set to 30 W/cm², as measured in water before skull transmission. TUS was delivered as bilateral sequential stimulation, with the transducer repositioned and realigned to the contralateral target using neuronavigation between hemispheres. In the GPi cohort, the left hemisphere was sonicated first, followed by the right [17]; in the STN cohort, the order of hemispheres was randomized equally across participants [16]. Individualized acoustic modelling and neuronavigation were used to guide transducer placement and optimize ultrasound delivery to M1 or GPi. Structural MRI scans were used to identify the target and determine the optimal sonication trajectory, accounting for skull geometry and individual anatomical differences. Acoustic and thermal simulations using BabelBrain were used to estimate the transmitted pressure distribution, focal location, and in-situ exposure after skull transmission. The three stimulation–recording configurations are illustrated in Fig. 1B. Patients were assessed during the ON-medication state, with DBS switched off during LFP recordings. In the STN cohort, active stimulation and sham were administered in separate sessions, whereas in the GPi cohort, sham and active sonication were delivered within the same visit, separated by a 30-minute washout interval. STN LFPs were recorded at rest before and during sonication and at 10, 30, and 45 minutes afterwards. Separate finger-tapping recordings were obtained at baseline and at post-sonication time points. In the participant with GPi recordings (sub08), resting LFPs were acquired before and during sonication and at 10, 25, and 40 min afterwards. No finger-tapping recordings were obtained in the GPi cohort.

### 2.4. FTG identification and Time–frequency visualization

Individual FTG peaks were identified from bipolar LFP recordings after separating narrowband oscillatory activity from the aperiodic spectral background (Fig. 1C). For each recording, power spectra were estimated using the multitaper method with 4-s windows, 50% overlap, and a time–bandwidth product of 4. Spectra were averaged across windows, and the aperiodic component was modelled as a linear fit in log-frequency versus log-power (dB) space over 35–95 Hz, excluding the 70–90 Hz FTG band. The fitted component was then subtracted from the power spectrum to obtain the residual spectrum. A candidate peak within 70–90 Hz was classified as FTG if its residual power exceeded the mean plus two standard deviations of the residual power in the 60–65 Hz reference band. Candidate peaks were visually inspected to confirm FTG detection. Participants were included in the longitudinal analysis if FTG was present on at least two bipolar contacts in the pre-sonication resting recording of an active sonication session, as a pre-sonication peak was required to define each contact’s FTG frequency and baseline. This procedure was applied to identify reliable narrowband peaks across recording contacts and sessions. For longitudinal analysis, resting and finger-tapping recordings were divided into non-overlapping 4-s windows. For each contact, FTG power was averaged within ±4 Hz of the pre-sonication peak frequency, restricted to the 70–90 Hz range. Power was expressed in dB relative to the 60–65 Hz reference-band power within the same window and subsequently referenced to the mean pre-sonication value of the corresponding contact. Thus, the reported FTG changes represent changes in normalized narrowband gamma power rather than absolute gamma-band power. Individual baseline peak frequencies are reported in Table 1.

**Table 1.** FTG peak frequencies in the pre-stimulation resting recordings.

| **Participant** | **Sonication target** | **Recording site** | **No. of contacts with FTG** | **Peak range (Hz)** |
|---|---|---|---|---|
| **sub09** | M1 | STN | 3 | 76.7–77.1 |
| **sub11** | M1 | STN | 3 | 77.7–78.4 |
| **sub12** | M1 | STN | 2 | 82.7–83.6 |
| **sub03** | GPi | STN | 2 | 83.3–83.9 |
| **sub08** | GPi | GPi | 3 | 74.4–74.6 |

### 2.5. Relationship of FTG to beta power, levodopa dose, and dyskinesia

Cross-sectional analyses included all 27 patients from the STN (n = 17) and GPi (n = 10) cohorts. All recordings were obtained during the ON-medication state, and levodopa-equivalent daily dose (LEDD) was obtained from clinical records. Patients were classified as FTG-positive if a confirmed peak was detected during at least one recording session and as FTG-negative otherwise. In the STN cohort, eight patients were FTG-positive and nine were FTG-negative. In the GPi cohort, one participant (sub08) was FTG-positive and nine were FTG-negative. Beta and FTG power were estimated from pre-sonication resting recordings using Welch’s method with 2-s Hann windows and 50% overlap. Relative beta power was defined as power in the 13–30 Hz band expressed as a percentage of the total power between 5 and 45 Hz. Relative FTG power was computed from the same aperiodic residual approach as in Section 2.4, taking the maximum 3-Hz-smoothed residual within 70–90 Hz after referencing to the 60–65 Hz band. For each measure, the contact with the highest value was selected within each session and hemisphere. Patient-level values were then obtained by averaging across sessions and hemispheres. LEDD was compared between FTG-positive and FTG-negative patients using a two-sided Mann–Whitney U test. Among FTG-positive patients, associations between relative FTG and relative beta power, and relative FTG and LEDD were assessed using Spearman’s rank correlation. The relationship between FTG and dyskinesia was examined at the session level in the STN cohort. Dyskinesia was assessed using the Movement Disorder Society–Unified Parkinson’s Disease Rating Scale (MDS-UPDRS III) before and after sonication. A session was classified as dyskinesia-positive if dyskinesia was present at either assessment and as FTG-positive if a confirmed FTG peak was detected in any resting recording from that session. The association between FTG and dyskinesia was examined across 47 sessions from 17 patients using mixed-effects logistic regression, with FTG as a fixed effect and participant as a random intercept (dyskinesia ∼ FTG + (1 | patient)) to account for repeated sessions within patients.

### 2.6. Spatial distribution of FTG within the STN

To examine where FTG was recorded relative to the STN, we used the contact coordinates of the STN cohort (Section 2.2; MNI ICBM 2009b nonlinear asymmetric space [16]). Each participant’s sensing configuration included six bipolar channels: contacts 0–3, 1–3 and 0–2 in each hemisphere. Each channel was placed at the midpoint between its two contact levels. Right-hemisphere channels were mirrored onto the left hemisphere, so all 102 channels (17 patients × 6) could be shown relative to a single STN. A channel was classified as FTG-positive if a confirmed peak (Section 2.4) was detected in at least one resting recording from any session. Channels included in the longitudinal analysis were additionally marked by the direction of their mean post-sonication FTG change. STN and motor-STN boundaries were taken from the DISTAL atlas [22], which is defined in the same template space, and projected onto coronal, sagittal and axial planes (Fig. 5A). To relate FTG to the dorsal border of the STN, we computed each channel’s height as its z coordinate minus the highest atlas STN voxel at the channel’s x–y position. Channels were then grouped by height (< −2, −2 to 0, 0 to 2 and > 2 mm), and the percentage of FTG-positive channels was computed for each group (Fig. 5B).

To test whether FTG presence depended on the spatial position of the recording channels, we fitted logistic regression models with FTG (present/absent) as the outcome and channel coordinates as predictors. Coordinates were centered on the centroid of the atlas STN. Candidate models included every combination of x, y and z main effects and their pairwise interactions, with each interaction included only alongside both of its main effects. Models were compared using the Akaike information criterion (AIC). Because each participant contributed six channels, observations were not independent, so we assessed the selected model in three further ways. First, standard errors were made cluster-robust, with patients as clusters. Second, a participant-level permutation test reassigned each participant’s six-channel FTG pattern as a block to another participant’s lead positions (5,000 permutations). Third, the model was refitted within FTG-positive patients only, to test location within leads independently of which patients expressed FTG. A mixed-effects model with participant as a random effect was also fitted, but the between-participant variance did not converge to a stable estimate, because nine of the 17 patients had no FTG on any channel.

### 2.7. Statistical analysis

Changes in FTG power relative to the pre-sonication baseline were assessed separately for each participant using linear mixed-effects models with recording epoch as a fixed effect and recording contact as a random intercept (FTG ∼ Epoch + (1 | Channel)). The pre-sonication epoch served as the reference condition. P-values for comparisons across epochs were adjusted within each participant using the Benjamini–Hochberg false discovery rate (FDR) procedure. For finger-tapping analyses, each 4-s task window was compared with the mean resting FTG power of the corresponding contact at the same recording epoch. The resulting task–rest differences were assessed separately for each participant and epoch using a linear mixed-effects model with contact as a random intercept (Difference ∼ 1 + (1 | Channel)). P-values were adjusted across the four assessed epochs (baseline and 10, 30, and 45 minutes after sonication) using the FDR procedure. For patients with analyzable sham sessions, active-versus-sham differences were assessed relative to the corresponding pre-sonication baseline within each session. The effects of stimulation condition, recording epoch, and their interaction were evaluated using a linear mixed-effects model with recording contact as a random intercept (FTG ∼ Condition × Epoch + (1 | Channel)). Differences between active and sham conditions were assessed through the Condition × Epoch interaction, with FDR correction across recording epochs. Two patients had sufficiently reproducible FTG during sham sessions to permit these comparisons. Sub09 contributed resting recordings from M1-sonication and passive-sham sessions, whereas sub11 contributed finger-tapping recordings from M1-sonication and active occipital-sham sessions. Unless otherwise specified, error bars represent the standard error of the mean (SEM) across recording windows rather than across patients. Statistical significance was defined as FDR-adjusted p < 0.05.

## 3. Results

### 3.1. FTG detection and baseline characteristics

In the STN cohort, detectable FTG was identified in eight of 17 patients during at least one recording session. Of these, four exhibited sufficiently reproducible FTG peaks across an active sonication session to permit longitudinal analysis: sub09, sub11, and sub12 during primary motor cortex (M1) sonication, and sub03 during GPi sonication. In the GPi cohort, reproducible FTG was detected in one of 10 patients (sub08). The longitudinal analysis therefore included five patients across three stimulation–recording configurations: M1 sonication with STN recordings (n = 3), GPi sonication with STN recordings (n = 1), and GPi sonication with GPi recordings (n = 1). All contacts included in the longitudinal analysis exhibited FTG during the pre-sonication resting recordings, with peak frequencies ranging from 76.7 to 83.9 Hz in the STN and 74.4 to 74.6 Hz in the GPi (Table 1).

### 3.2. M1 sonication with STN recordings

M1 sonication was associated with reduced resting STN FTG power in all three evaluable patients, although the magnitude and temporal profile of the response varied (Figs. 2A–C and 3A, left). In sub09, FTG power decreased by 4.8 dB during sonication (change from baseline: −4.8 dB, 95% CI −5.7 to −3.8), with reductions persisting at 10 minutes (−3.1 dB, 95% CI −3.8 to −2.3), 30 minutes (−2.0 dB, 95% CI −2.7 to −1.2), and 45 minutes (−2.5 dB, 95% CI −3.3 to −1.8; all adjusted P < 0.001). The corresponding spectrogram showed a prominent narrowband peak before sonication that became weaker and less continuous during subsequent recording epochs (Fig. 2A). In sub11, FTG decreased during sonication (−0.8 dB, 95% CI −1.1 to −0.4; P < 0.001), but did not differ significantly from baseline at 10 min (−0.2 dB, 95% CI −0.5 to 0.1; P = 0.14). FTG was again reduced at 30 min (−0.9 dB, 95% CI −1.2 to −0.7) and 45 minutes (−0.6 dB, 95% CI −0.9 to −0.3; both P < 0.001). The corresponding spectrogram showed variations in the magnitude and persistence of FTG suppression (Fig. 2B). In sub12, FTG power decreased more modestly but remained consistently below baseline during and after sonication, with reductions ranging from −0.7 to −1.1 dB across recording epochs (adjusted P < 0.001 at each epoch; Figs. 2C and 3A, left).

**Figure 2.**
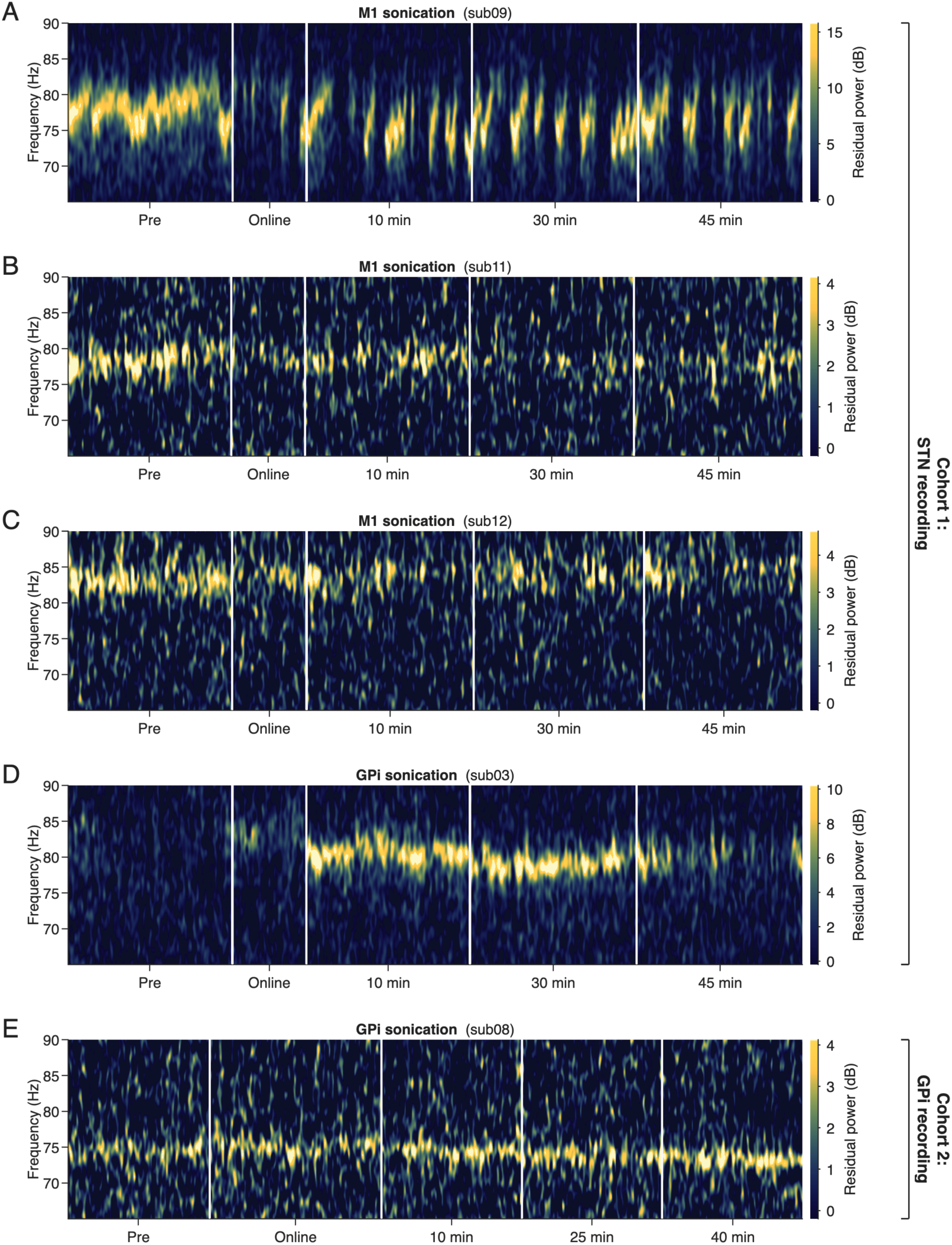
Individual residual-power FTG spectrograms. Panels A–D are from cohort 1 (STN recording) and panel E from cohort 2 (GPi recording). STN recordings after M1 sonication in (A) sub09, (B) sub11 and (C) sub12, STN recording after GPi sonication in (D) sub03, and GPi recording after GPi sonication in (E) sub08, shown before, during (online) and after sonication; Post-sonication recordings were obtained at 10, 30 and 45 minutes for A–D and at 10, 25 and 40 min for E.

**Figure 3.**
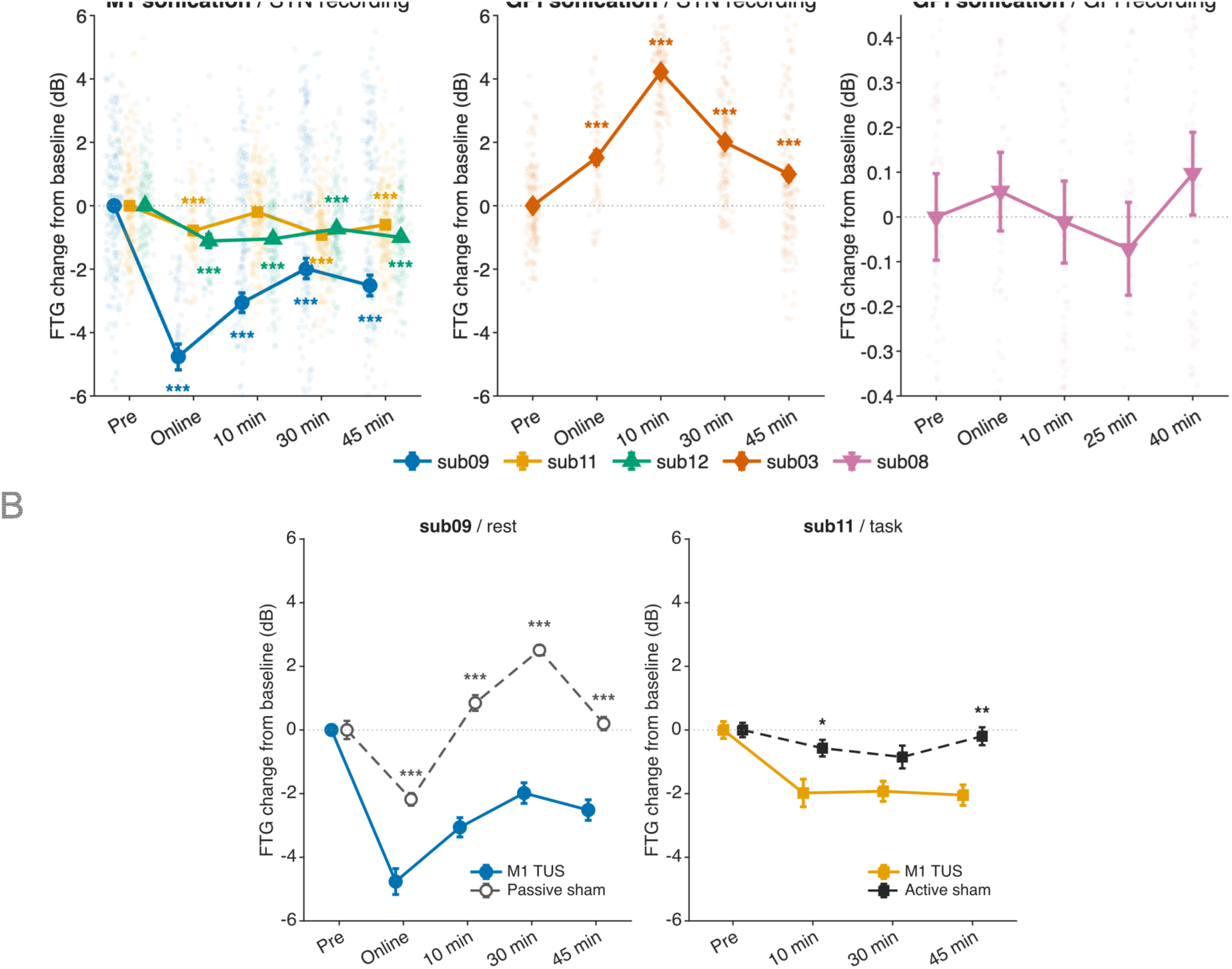
FTG changes after M1 and GPi sonication and comparison with sham. **(A)** Within-participant FTG changes relative to the pre-sonication baseline at rest are shown. M1 sonication decreased STN FTG (sub09, sub11, sub12) (left), whereas GPi sonication increased STN FTG (sub03) (middle) but produced no significant change in locally recorded GPi FTG (sub08) (right). Dots represent individual 4 s windows; markers and error bars indicate mean ± SEM across windows. Asterisks indicate post-sonication time points that differ from baseline in a linear mixed-effects model fitted per participant (FTG ∼ Epoch + (1 | Channel)). **(B)** FTG change after M1 sonication compared with sham in the two patients with FTG present in both sessions: sub09 at rest, compared with passive sham, and sub11 during finger tapping, compared with active sham. In both patients, M1 sonication resulted in greater FTG suppression compared with sham stimulation. Each session is referenced to its own pre-sonication baseline. Asterisks indicate time points at which the change after M1 sonication differed from the change after sham (Condition × Epoch interaction in a linear mixed-effects model, FTG ∼ Condition × Epoch + (1 | Channel)). In A and B, p-values were adjusted across time points within each comparison using the false discovery rate procedure (*p < 0.05, **p < 0.01, ***p < 0.001).

### 3.3. GPi sonication with STN or GPi recordings

In contrast to M1 sonication, GPi-targeted sonication was followed by a pronounced increase in resting STN FTG power in sub03 (Figs. 2D and 3A, middle). FTG increased during sonication (+1.5 dB relative to baseline, 95% CI 1.0 to 2.0) and reached its maximum at 10 min (+4.2 dB, 95% CI 3.8 to 4.6). The increase gradually declined but remained above baseline at 30 min (+2.0 dB, 95% CI 1.6 to 2.4) and 45 minutes (+1.0 dB, 95% CI 0.6 to 1.4; all adjusted P < 0.001). The corresponding STN spectrogram showed a narrowband oscillation near 80 Hz that became more prominent after sonication and gradually weakened during subsequent recording epochs (Fig. 2D).

GPi-targeted sonication did not significantly change locally recorded FTG in the GPi in sub08, despite the presence of a reproducible baseline peak (Figs. 2E and 3A, right). Relative to baseline, FTG power changed by +0.06 dB during sonication, −0.01 dB at 10 min, −0.07 dB at 25 min, and +0.10 dB at 40 min (reported 95% CI half-widths ≤0.27 dB; all adjusted P ≥ 0.88). The corresponding spectrogram showed a stable narrowband peak near 74 Hz across recording epochs (Fig. 2E).

### 3.4. M1 sonication versus sham

Two patients exhibited sufficiently reproducible FTG during their respective sham sessions, allowing within-participant comparisons between active M1 sonication and sham stimulation (Fig. 3B). In sub09, resting STN FTG decreased more following M1 sonication than during passive sham across all sampled epochs. Active-minus-sham differences in FTG change from the corresponding session baseline were −2.6 dB during sonication, −3.9 dB at 10 min, −4.5 dB at 30 min, and −2.7 dB at 45 minutes (all adjusted P < 0.001). During passive sham, FTG decreased transiently by 2.2 dB during the sham period before returning toward or above baseline, with changes of +0.9 dB at 10 min, +2.5 dB at 30 min, and +0.2 dB at 45 minutes. In contrast, FTG remained below baseline throughout the post-sonication recordings following active M1 stimulation. In sub11, FTG power recorded during finger tapping decreased by approximately 2 dB following M1 sonication, whereas changes following active occipital sham ranged from −0.2 to −0.9 dB. Relative to sham, M1 sonication produced greater reductions at 10 min (−1.4 dB, adjusted P = 0.034) and 45 minutes (−1.9 dB, adjusted P = 0.009). The difference at 30 min did not reach significance (−1.1 dB, adjusted P = 0.080). No other patients had sham-session recordings with sufficiently reproducible FTG on corresponding contacts to permit matched comparisons.

### 3.5. Relationship of FTG to beta power, levodopa dose, and dyskinesia

We examined the relationship between FTG expression and resting beta activity, levodopa-equivalent daily dose (LEDD), and dyskinesia (Fig. 4). Among FTG-positive patients (n = 9), relative FTG power showed nonsignificant correlation with relative beta power (Spearman’s ρ = 0.58, P = 0.11; Fig. 4A). LEDD did not differ significantly between FTG-positive and FTG-negative patients (median 1270 versus 1482 mg/day, respectively; n = 9 versus 18; P = 0.43; Fig. 4B). Within the FTG-positive group, relative FTG power was not significantly correlated with LEDD (ρ = 0.37, P = 0.34; Fig. 4B). We further examined the relationship between FTG detection and dyskinesia across 47 sessions from the 17 patients in the STN cohort. Dyskinesia was present in 8 of 17 sessions with detectable FTG (47%), compared with 6 of 30 sessions without FTG (20%; Fig. 4C). Although dyskinesia was more frequent during FTG-positive sessions, the association did not reach statistical significance in the mixed-effects logistic regression accounting for repeated sessions within patients (P = 0.067).

**Figure 4.**
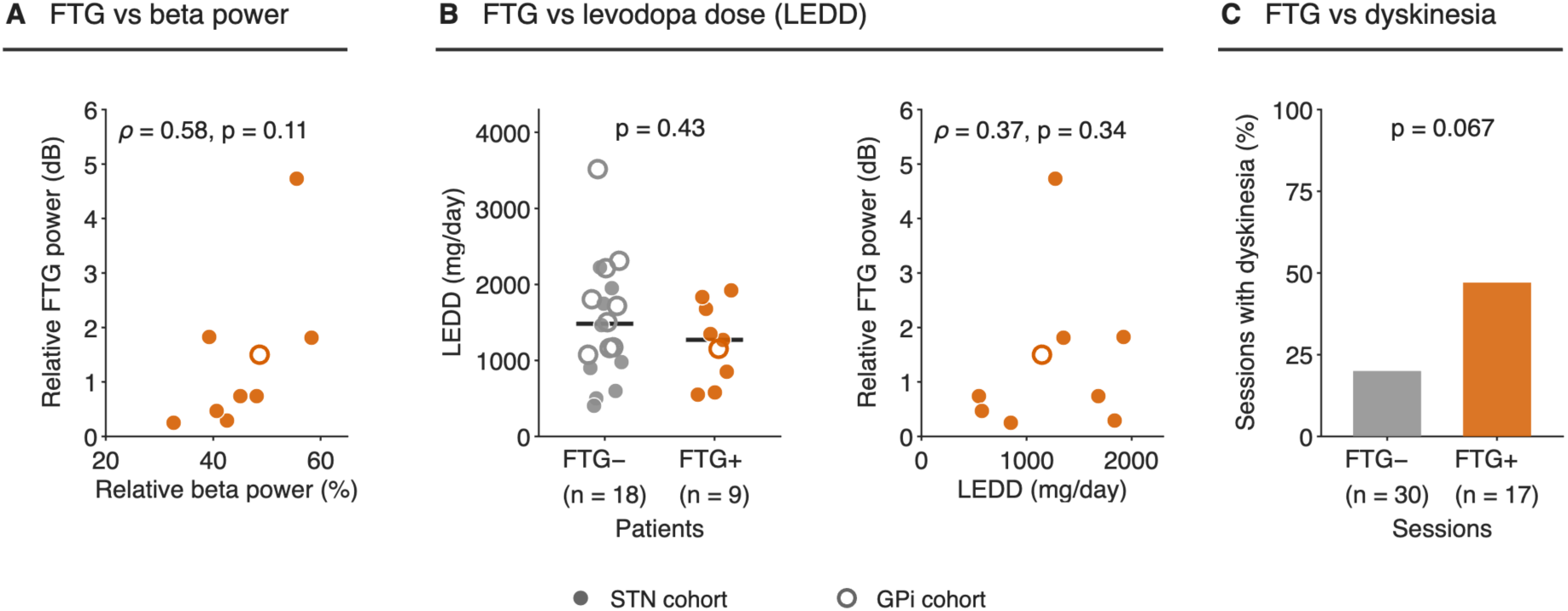
Relation of finely tuned gamma to beta power, levodopa dose and dyskinesia. Each dot is one patient: STN-cohort patients are filled, and GPi-cohort patients are hollow (FTG-positive, orange; FTG-negative, grey). All recordings were made on dopaminergic medication. Beta and FTG measures come from the pre-sonication rest recordings **(A)** Relative FTG power against relative beta power in FTG-positive patients (n = 9: eight STN-cohort patients and one GPi-cohort patient). **(B)** Left: levodopa-equivalent daily dose (LEDD) in FTG-negative (n = 18: nine STN-cohort and nine GPi-cohort patients) and FTG-positive (n = 9) patients; horizontal bars are medians, and p is from a two-sided Mann–Whitney U test. Right: LEDD against relative gamma power in FTG-positive patients. **(C)** Percentage of sessions with dyskinesia (present on the pre- or post-sonication MDS-UPDRS III rating) among sessions without FTG (6 of 30 sessions, 20%) and with FTG (8 of 17 sessions, 47%), across 47 sessions from 17 STN-cohort patients. p is from a mixed-effects logistic regression (dyskinesia ∼ FTG + (1 | patient)).

### 3.6. Finger-tapping modulation of STN FTG

Finger tapping produced heterogeneous changes in STN FTG power across patients, both before and after sonication (Supplementary Fig. S1). At baseline, FTG power was higher during finger tapping than at rest in sub09 (+2.1 dB, 95% CI 1.6 to 2.6) and sub11 (+1.7 dB, 95% CI 1.1 to 2.2), whereas it was reduced in sub12 (−1.8 dB, 95% CI −2.5 to −1.1; all adjusted P < 0.001). In sub03, finger tapping produced no significant change relative to rest (+0.5 dB, 95% CI −0.2 to 1.3; P = 0.17). Following M1 sonication, sub09 continued to exhibit task-related increases in FTG power at 10 min (+3.2 dB) and 45 minutes (+4.0 dB; both adjusted P < 0.001). The increase at 30 min did not reach statistical significance (+1.6 dB, P = 0.053). In sub11, task-related changes were no longer significant at any post-sonication epoch, with the largest difference observed at 30 min (+0.7 dB, P = 0.053). In sub12, the task-related reduction remained significant only at 30 min (−1.2 dB, P < 0.001). Following GPi sonication in sub03, FTG power during finger tapping was higher than at rest by +1.8 dB at 10 min (P = 0.17) and +1.0 dB at 30 min (P = 0.098), although neither difference reached significance. At 45 minutes, the task-related increase was more pronounced (+3.4 dB, 95% CI 2.5 to 4.3; P < 0.001). All reported P-values for the task comparisons were adjusted across the four tested epochs within each participant using the FDR procedure.

### 3.7. Spatial distribution of FTG within the STN

Across the STN cohort, FTG was detected on 35 of 102 bipolar channels. FTG-positive channels were located both within and dorsal to the STN (Fig. 5A). Channels included in the longitudinal analysis lay mostly in the dorsal STN and the region above it; the two channels with increased FTG after GPi sonication (sub03) were within the dorsolateral STN. The proportion of channels with FTG increased with height relative to the dorsal STN border: 19% (6 of 31) of channels more than 2 mm inside the border, 38% (17 of 45) between −2 and 0 mm, 45% (9 of 20) between 0 and 2 mm, and 50% (3 of 6) more than 2 mm above it (Fig. 5B). In logistic regression, the best-fitting model included anteroposterior (y) and dorsoventral (z) position and their interaction (ΔAIC = 2.0 relative to the next-best model, which additionally included mediolateral position). In this model, FTG was more likely at more dorsal and more posterior positions (likelihood-ratio χ²(3) = 17.2, P < 0.001; McFadden R² = 0.13). However, the effect was sensitive to within-participant clustering. With participant-clustered standard errors, only the y × z interaction remained significant (P = 0.029). Neither the participant-level permutation test (P = 0.073) nor the model fitted within the eight FTG-positive patients (48 channels; P = 0.11) reached significance. The spatial pattern may therefore partly reflect which patients expressed FTG rather than position within individual leads and should be considered preliminary.

**Figure 5.**
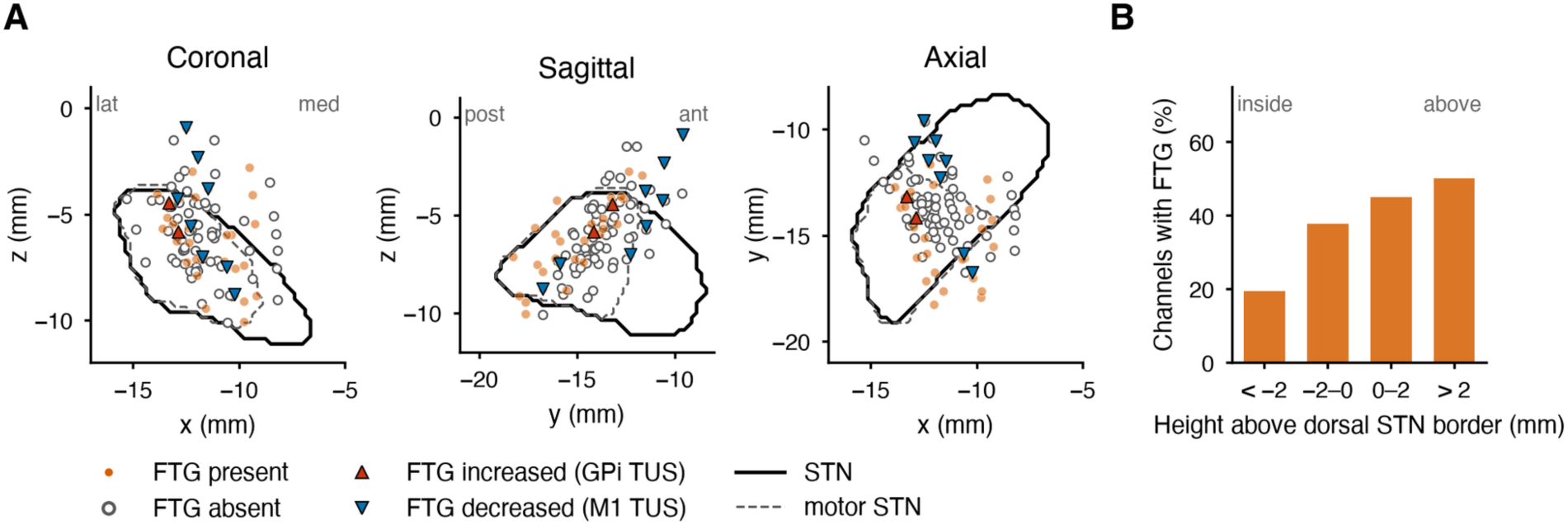
Spatial distribution of FTG across STN recording channels. (A) Midpoints of all bipolar recording channels from the STN cohort (17 patients × 6 channels = 102) in MNI space, shown in coronal, sagittal and axial projections, with right-hemisphere channels mirrored onto the left. Filled orange dots are channels with FTG detected in at least one resting recording (n = 35); open grey circles are channels without detectable FTG (n = 67). Triangles mark channels in the longitudinal analysis, coloured by the direction of their mean post-sonication FTG change: red upward triangles, increase after GPi sonication (sub03); blue downward triangles, decrease after M1 sonication (sub09, sub11 and sub12). Solid and dashed outlines show the STN and its motor subregion from the DISTAL atlas. (B) Percentage of channels with FTG by height relative to the dorsal STN border (negative, inside the STN; positive, above it).

## 4. Discussion

The present study provides preliminary evidence that endogenous FTG can be modulated by noninvasive TUS of cortical and subcortical motor-network targets. M1 sonication was associated with reduced FTG power recorded from the STN, whereas GPi sonication increased STN FTG in one participant but did not significantly alter locally recorded GPi FTG in another. Importantly, resting FTG was not significantly correlated with beta-band activity or levodopa-equivalent daily dose (LEDD), and medication dosage did not differ between FTG-positive and FTG-negative patients. In contrast, dyskinesia was more frequently observed in sessions with detectable FTG, although this association did not reach statistical significance. Together, these observations suggest that FTG may capture aspects of motor-network physiology not fully reflected by beta activity or medication dosage, while retaining a potential relationship with dyskinesia. These findings support further investigation of FTG as a complementary electrophysiological biomarker for neuromodulation in Parkinson’s disease.

### 4.1. Target-dependent and circuit-dependent modulation of FTG

The contrasting FTG responses following M1 and GPi sonication suggest that ultrasound-induced modulation may be shaped by the functional organization of cortico–basal ganglia circuits rather than reflecting a uniform excitatory or inhibitory effect. M1 influences STN activity directly through the hyperdirect pathway and indirectly through broader corticostriatal and basal ganglia circuitry [23–25]. Accordingly, reduced STN FTG following M1 sonication may reflect altered cortical drive, cortico-subthalamic coupling, or the network’s propensity to express narrowband gamma oscillations. Conversely, GPi occupies a major output position within the basal ganglia and participates in interconnected pallido-thalamo-cortical and subthalamo-pallidal circuits [24–26]. Increased STN FTG following GPi sonication may therefore reflect changes in network coupling or oscillatory dynamics rather than a direct increase in local neuronal excitability. Recent evidence of endogenous FTG within the GPi [27], and the finding that electrical stimulation of either the STN or the GPi entrains cortical narrowband gamma [12], further support the view that FTG reflects distributed motor-network activity. The contrasting STN responses to M1 and GPi sonication suggest that its expression also depends on which node is stimulated and on the physiological context.

GPi-targeted sonication did not significantly alter locally recorded FTG in the single evaluable participant with GPi leads, despite a reproducible baseline peak. This difference may reflect the distinct physiological sensitivity of local GPi activity compared with STN responses, as well as differences in electrode location and spatial sampling of oscillatory activity. Direct sonication near implanted electrodes may also introduce recording artifacts during stimulation, potentially increasing variability in the measured LFP signal [28, 29]. However, such artifacts would not readily explain the absence of changes in subsequent post-sonication recordings.

The sham observations provide additional, although limited, support for the target dependence of these effects. The parent studies incorporated control conditions, including passive sham and stimulation of the occipital cortex as an active sham [16, 17], but reliable FTG was not consistently detectable across sham recordings. Only two patients exhibited sufficiently reproducible FTG during sham sessions to permit matched comparisons with active M1 sonication. In these patients, FTG suppression following M1 stimulation was greater than during sham at selected recording epochs. The limited detectability of FTG during control sessions may reflect its intermittent expression across recording conditions [5].

### 4.2. Regional differences in FTG detection across the basal ganglia

An additional observation was the relatively low prevalence of consistently detectable FTG across the two recording cohorts. Among 17 patients with STN leads, eight exhibited FTG during at least one recording session, whereas four (24%) showed sufficiently reproducible peaks across the relevant stimulation session for longitudinal analysis. In contrast, reproducible FTG was identified in only one of 10 patients (10%) with GPi leads. These observations are broadly consistent with previous studies reporting heterogeneous and intermittent FTG expression across patients and recording sites. In a recent study involving 67 patients with STN DBS, spontaneous FTG was detected in 19% of recorded STNs [5]. Although recording conditions, detection criteria, and denominators differed, the proportion of patients with consistently detectable STN FTG in our study was of a similar order, particularly given the requirement for reproducible peaks across repeated recording epochs.

The lower detection rate in the GPi cohort raises questions about the regional organization and expression of gamma oscillations within basal ganglia circuits. The STN receives direct excitatory input from cortical motor regions and participates in recurrent interactions with the external globus pallidus, providing a potential substrate for synchronized oscillatory activity [23, 26, 30]. In contrast, GPi integrates excitatory inputs from the STN with inhibitory inputs from the striatum and external globus pallidus, which may influence the local synchronization and expression of gamma activity [24, 25]. Early simultaneous recordings from STN and GPi demonstrated levodopa-induced oscillations near 70 Hz in both nuclei, with coherent activity and STN-leading-GPi phase relationships [6, 8]. These findings suggest that FTG recorded within the GPi may, at least partly, reflect oscillatory activity transmitted through interconnected basal ganglia circuits. Narrowband gamma has been proposed to arise from recurrent excitatory–inhibitory interactions within cortico–basal ganglia circuits, with cortical gamma often prominent and coherent with subcortical sites [2, 7, 8]. Computational work further suggests that STN stimulation can engage gamma-range dynamics in striatal fast-spiking interneuron networks [32], consistent with the general role of local inhibitory interactions in gamma generation [31]. However, the available evidence does not establish distinct anatomical sources of FTG or whether its generation differs systematically between STN and GPi [27]. Larger studies incorporating standardized recording conditions, simultaneous STN and GPi recordings, and detailed electrode localization are needed to determine whether FTG detectability differs systematically between these nuclei.

Electrode location may influence FTG detection. The proportion of bipolar channels with detectable FTG increased with height relative to the atlas-defined dorsal STN border (Fig. 5). Previous intraoperative recordings identified stronger gamma activity near the upper STN and bordering zona incerta, where neuronal discharges were locked to the gamma oscillation [33]. Our spatial pattern may therefore partly reflect differences in how closely recording contacts sample neuronal populations participating in gamma synchronisation. The physiological relevance of this activity is supported by simultaneous recordings showing that periods of stronger gamma were accompanied by higher average neuronal firing rates and greater potential information-coding capacity in the subthalamic region [34].

### 4.3. Modulation of endogenous FTG versus gamma entrainment

Our observations reflect modulation of endogenous FTG power rather than demonstrated gamma entrainment. High-frequency DBS can entrain gamma oscillations at approximately half the stimulation frequency [11, 12], with evidence linking this activity to motor improvement [11] and the therapeutic ON state [5]. Similar subharmonic entrainment has recently been demonstrated during thalamic DBS for multiple sclerosis tremor [14]. Together with earlier reports of endogenous narrowband thalamic gamma [13], this suggests that the capacity for gamma synchronization extends beyond the parkinsonian basal ganglia. In contrast, the present theta-burst TUS protocol altered the amplitude of pre-existing FTG without establishing evidence of stimulation-locked gamma activity.

This distinction may be important for understanding how electrical and ultrasound stimulation interact with ongoing neural oscillations. Electrical DBS delivers repetitive stimulation directly to neural tissue and can entrain ongoing activity through frequency-dependent interactions with oscillatory networks [11, 12]. TUS, conversely, may influence the expression of endogenous rhythms through changes in neuronal excitability and synaptic interactions without necessarily imposing a new oscillatory frequency [15]. The persistence of FTG changes beyond sonication suggests that the observed effects were not restricted to the period of acoustic stimulation. However, these recordings cannot establish the underlying cellular mechanisms or determine whether sustained responses reflect changes in synaptic efficacy.

The movement-related findings further highlight the context-dependent expression of FTG. During finger tapping, STN FTG increased relative to rest in two patients, decreased in one, and showed no detectable baseline change in another. These responses also varied following sonication, including the emergence of a significant task-related increase after GPi stimulation in one participant. Previous studies have demonstrated movement-related changes in endogenous FTG [10], while DBS-entrained gamma has been shown to increase during repetitive finger tapping and to be associated with improved motor performance [11]. The heterogeneous changes observed here suggest that endogenous FTG does not necessarily increase uniformly during motor execution but may depend on individual network states and the interaction between movement and stimulation. These observations further support the importance of considering behavioral state when interpreting FTG as an electrophysiological marker.

### 4.4. Relationship to beta activity, dopaminergic medication, dyskinesia, and biomarker relevance

An important finding was the absence of a statistically significant correlation between resting FTG and beta-band power (ρ = 0.58, P = 0.11; Fig. 4A). In our STN cohort, M1- and GPi-targeted TUS produced contrasting effects on STN beta oscillations, with motor improvement observed following M1 stimulation [16]. The present analysis extends these observations to narrowband gamma and suggests that FTG and beta activity may provide complementary information about basal ganglia network states. Beta oscillations are predominantly associated with the hypokinetic state in PD, whereas FTG has been linked to dopaminergic ON states, movement, and, under certain conditions, dyskinesia [1–5, 9]. Although the positive FTG–beta correlation did not reach statistical significance, this does not establish that the two rhythms are independent. Their different physiological associations nevertheless suggest that FTG may capture aspects of motor-network activity not fully reflected by beta oscillations alone.

Previous electrophysiological studies have also demonstrated interactions between oscillatory frequency bands in the basal ganglia. López-Azcárate et al. [35] reported coupling between beta-band phase and the amplitude of high-frequency oscillations in the STN, suggesting that interactions across frequency bands may contribute to abnormal network synchronization in PD. Although the high-frequency oscillations examined in that study differ from the 65–90 Hz FTG investigated here, these findings highlight the importance of considering relationships between neural rhythms rather than interpreting each frequency band in isolation. More recently, spontaneous and stimulation-entrained FTG have been shown to coexist with beta activity, with their relationships varying according to stimulation and dopaminergic state [5]. Thus, FTG and beta oscillations may reflect overlapping but potentially complementary aspects of motor-network physiology, supporting their combined investigation as electrophysiological biomarkers.

We also found no significant relationship between FTG and LEDD (Fig. 4B). Medication dosage did not differ significantly between FTG-positive and FTG-negative patients, and resting FTG power was not significantly correlated with LEDD among FTG-positive patients. These observations contrast with our previous GPi-TUS study, in which higher dopaminergic medication doses were associated with greater ultrasound-induced changes in total LFP power [17]. The present findings therefore suggest that endogenous FTG expression does not necessarily follow the same medication-dose relationship observed for broader electrophysiological responses. This distinction is particularly relevant because FTG has been shown to track medication cycles and dopaminergic ON states [4, 5]. However, all recordings in the present study were obtained during the ON-medication state, and the lack of a significant relationship with LEDD does not exclude sensitivity to medication timing or changes in dopaminergic state.

In contrast to the lack of significant associations with beta activity and LEDD, FTG showed a trend toward an association with dyskinesia. Dyskinesia was present in 8 of 17 sessions with detectable FTG (47%), compared with 6 of 30 sessions without FTG (20%; Fig. 4C). Although this association did not reach statistical significance after accounting for repeated sessions within patients (P = 0.067), the direction of the effect is consistent with earlier studies linking narrowband gamma activity to levodopa-induced dyskinesia [1, 2]. This observation suggests that dyskinesia may be a relevant clinical correlate of FTG expression in our cohort. Nevertheless, recent investigations have demonstrated that FTG can also occur during therapeutic ON states without accompanying dyskinesia [4, 5]. Therefore, our findings are compatible with an association between FTG and dyskinesia but do not establish FTG as a dyskinesia-specific biomarker.

Together, these findings suggest that FTG may provide complementary information to beta oscillations for characterizing motor-network states in PD. The absence of significant associations with resting beta activity and LEDD, together with the observed trend toward greater dyskinesia prevalence during FTG-positive sessions, supports further investigation of FTG as an additional electrophysiological biomarker. However, the clinical significance of FTG modulation and its potential value for adaptive DBS require prospective validation.

### 4.5. Limitations and future directions

Several limitations should be considered. Although the parent studies included 17 patients with STN leads and 10 patients with GPi leads, the longitudinal analysis was restricted to five patients with sufficiently reproducible FTG. While the detection rate was broadly consistent with previous reports, the small sample limited systematic comparisons across stimulation targets, recording sites, and behavioural states. The cross-sectional analyses included 27 patients, but correlations involving FTG power were restricted to nine FTG-positive patients, limiting statistical power. Similarly, the observed association between FTG and dyskinesia was based on repeated sessions within a relatively small cohort and did not reach statistical significance. Moreover, only two patients had reliable FTG during matched sham recordings, limiting the ability to establish ultrasound-specific effects across the broader cohort. Differences in electrode location, recording sensitivity, and potential sonication-related artifacts may also have affected the interpretation of local FTG responses.

Future TUS studies incorporating quantitative assessments of bradykinesia and dyskinesia at multiple time points would be particularly informative. Longitudinal recordings could also clarify the reliability of FTG detection and whether individual FTG responses predict clinical or electrophysiological effects of ultrasound stimulation.

Overall, these preliminary findings suggest that endogenous FTG is sensitive to noninvasive perturbation of cortico–basal ganglia networks and may provide complementary information about motor-network states alongside beta oscillations. Although its clinical significance remains to be established, combining noninvasive TUS with intracranial electrophysiological sensing may offer a useful approach for investigating FTG as a candidate biomarker and its potential relevance for future adaptive DBS and individualized neuromodulation in PD.

