## Supplementary Figure S1 for "Target-dependent modulation of finely tuned gamma by transcranial ultrasound in Parkinson’s disease"

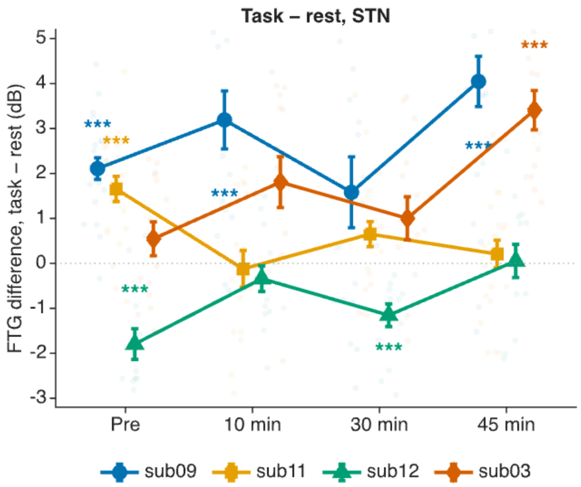


**Supplementary Figure S1. Movement-related modulation of STN FTG.** Difference between finger-tapping and resting normalized FTG power in M1-sonicated sub09, sub11 and sub12 and GPi-sonicated sub03, measured before sonication and at 10, 30 and 45 minutes after stimulation. Values indicate the mean task-minus-rest difference for the same contact and epoch; markers and error bars are mean ± SEM across 4-s task windows. Asterisks denote FDR-adjusted within-participant task–rest contrasts (*P < 0.05, **P < 0.01, ***P < 0.001).
